# An artificial nervous system for communication between wearable and implantable therapeutics

**DOI:** 10.1101/2025.06.04.657863

**Authors:** Ramy Ghanim, Yoon Jae Lee, Garan Byun, Joy Jackson, Julia Z. Ding, Elaine Feller, Eugene Kim, Dilay Aygun, Anika Kaushik, Alaz Cig, Jihoon Park, Sean Healy, Camille E. Cunin, Aristide Gumyusenge, Woon Hong Yeo, Alex Abramson

## Abstract

Bioelectronics have transformed our capacity to monitor and treat diseases; however, a lack of micrometer-scale, energy efficient communication options limit these devices from forming integrated networks that enable full-body, sensor driven, physiological control. Inspired by our nervous system’s ability to transmit information via ionic conduction, we engineered a Smart Wireless Artificial Nervous System (SWANS) that utilizes the body’s own tissue to transmit signals between wearables and implantables. When SWANS emits signals, it generates voltage gradients throughout the body that selectively turn on implanted transistor switches when exceeding their gate threshold voltages. SWANS’ implantable communication components maintain syringe-injectable footprints and >15x greater power efficiencies than Bluetooth and Near Field Communication. In vivo studies in rats demonstrate SWANS’ ability to wirelessly regulate dual hind leg motor control by connecting electronic-skin sensors to implantable neural interfaces via ionic signaling as well as coordinate bioelectronics throughout the epidermal, subcutaneous, intraperitoneal, and gastrointestinal spaces.

## Introduction

Wearable (*1, 2*), implantable (*3–6*), and ingestible (*7, 8*) bioelectronics monitor and treat the world’s most prevalent illnesses including diabetes (*9, 10*), cancer (*11, 12*), and heart disease (*13, 14*); however, almost all bioelectronics work independently from one another, limiting the capacity for biosensors to interact and remotely trigger therapeutics. In-body networks of electronic therapeutics are inherently limited by the large power and size requirements necessary for communication that limit implant battery lifetimes to weeks and necessitate invasive surgical implantation (*15–17*). One example of sensors and therapeutics working side-by-side is the wearable continuous glucose monitor and insulin pump; yet insulin pumps’ batteries, measuring 8 cm^3^, only last 7 days partially due to constant Bluetooth Low Energy (BLE) connectivity (*18*). Other wireless communication and power transfer methods, such as near-field communication (NFC), require bulky external antennae that limit patient adherence and require centimeter-scale alignment with targeted devices (*19, 20*). Here, we harness the conductive nature of body tissue to transmit communication signals ionically between wearable sensors and implantable devices. In the process, we create an artificial nervous system that: (1) reduces communication components to an emitted square-wave pulse and a single micrometer-scale transistor per implanted device; (2) independently triggers multiple devices throughout the tissue from a multiplexed wearable hub; and (3) extends the battery life of implanted devices compared to NFC and Bluetooth by 15x and 30x, respectively.

Here, we introduce SWANS, the Smart Wearable Artificial Nervous System. SWANS unlocks the potential for long-term closed-loop networks of wearable and in-body therapeutics by providing a framework for communication between dispersed bioelectronics via intertissue ionic conduction **(Fig. 1A)**. To enable effective communication, SWANS’ wearable biosensor hub emits electrical pulses that produce electric fields inside the body; then, the implantable receiving network detects the generated voltage gradient through tissue-interfacing pads to turn on a transistor switch which briefly enables a therapeutic action **(Fig. 1B)**. Each implant possesses a unique transistor circuit that only responds to its corresponding signal. The U.S. Food and Drug Administration has already confirmed the safety and efficacy of in-body ionic communication in humans for other uses (*21*); for example, Abilify MyCite pills utilize ionic conduction to transmit ingestion event data from an oral drug tablet to a wearable device by recording the presence or absence of an electrical signal in the body (*22, 23*). Preclinical studies have also utilized ionic conductance to similarly transmit data across tissues from implantable (*24*), wearable (*25*), and ingestible (*26*) sensors. Additionally, engineers have previously attempted to create body-networks of wearable sensors without including implants (*27, 28*); however, a fully functional body-therapeutic-network requires integration of both sensors and actuators across all tissue depths to enable closed-loop therapies. We demonstrate in vivo in rats that SWANS can wirelessly trigger subcutaneously implanted, intraperitoneally implanted, and gastrointestinally localized actuators via sensors and electronics placed on the epidermis. Furthermore, we demonstrate in vivo proof-of-concept by utilizing the SWANS architecture to wirelessly trigger implantable sciatic nerve stimulators for dual hind-leg animation. By detecting front limb motion and activating hind leg motion, we show that SWANS mimics how nervous systems selectively trigger and transmit targeted signals from one end of the body to another.

**Fig. 1.**
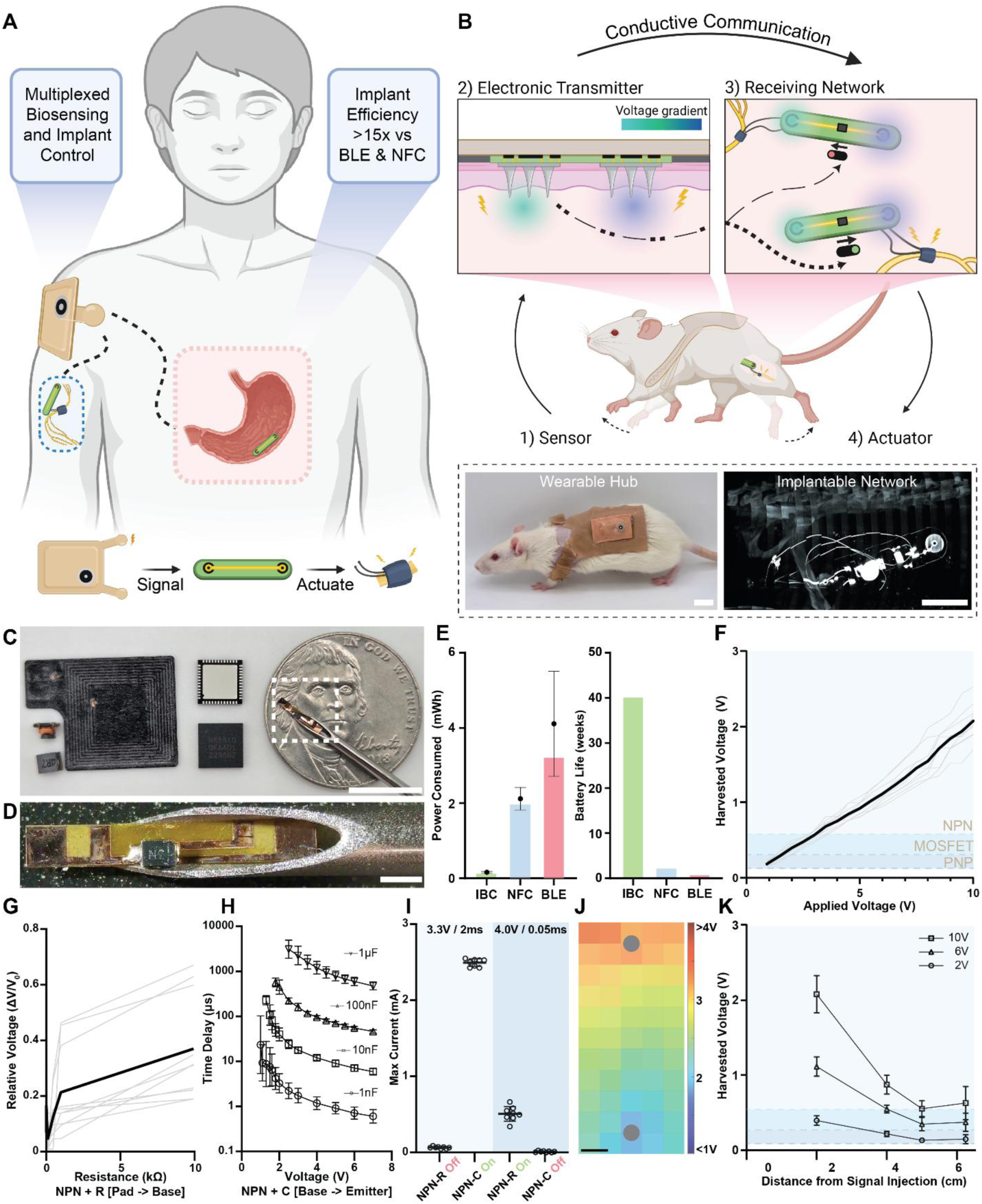
In-body communication enables low-power networks of wearable and implantable therapeutics. **(A)** A smart wireless artificial nervous system [SWANS] utilizes a wearable hub to control multiple therapeutic actuators throughout the subcutaneous, intraperitoneal, and gastrointestinal spaces by sending electrical signals ionically through tissue. **(B)** An example SWANS use case, where wearable sensors detect front limb motion, triggering a wearable hub to generate an electric field throughout the body. Distal implants receive the voltage gradient using tissue interfacing pads and turn on corresponding transistor switches. If the signals match, sciatic nerve stimulators are actuated for hind leg motor control. **(C)** Right to left: SWANS communication component; Bluetooth System-on-chip; millimeter-scale and centimeter-scale working distance NFC antennas. **(D)** Zoomed in SWANS communication component in 16-gauge needle. **(E)** Theoretical (bars) and experimental (points, Avg ± SD, n=3) power characteristics of implanted neuromodulation bioelectronics controlled via in-body ionic communication (IBC), Near Field Communication (NFC), and Bluetooth Low Energy (BLE). **(F)** Voltage measured across a transistor’s gate when applying increasing amplitude pulses across ex vivo chicken breast tissue. Shaded regions denote transistor gate and threshold voltages. (black = avg, grey = ind; n = 9) **(G)** Adding a resistor between the harvesting pad and NPN transistor base (NPN-R) ex vivo in chicken breast increases the required applied voltage for switching (black = avg, grey = ind; n = 10). **(H)** Adding a capacitor between the base and emitter of an NPN transistor (NPN-C) ex vivo in chicken breast increases the required applied pulse length for switching (Avg ± SD; n=6-11). **(I)** Low voltage long pulses only trigger NPN + capacitor switches ex vivo in chicken breast while high voltage short pulses only trigger NPN + resistor switches. (R=2.2 kΩ; C= 1 µF; ind points, Avg ± SD, n=8). **(J)** Heat map of generated voltage potentials upon application of [6 V, 1 Hz, 200 µs] pulses across two hypodermic needles (gray circles) in ex vivo chicken breast models (n=5). **(K)** Voltage measured across each transistor’s gate when moving the receiving pads away from the signal injection source. Measurements taken at different pulse amplitudes. (line = Avg ± SD, n = 9). Scale Bars: B, C, K = 1 cm; D = 1 mm.

### In-Body ionic communication for triggering bioelectronic therapeutics

SWANS shrinks the size of implanted communication components **(Fig. 1C-D)** and improves the battery life of implantable devices compared to those employing BLE and NFC communication by utilizing micrometer-scale, low-power transistor switches to regulate DC output and on/off states; this compares to BLE system on a chip components that measure at least 6 mm in width and NFC antennas that can reach centimeter scale in size. SWANS ensures that the current through the device remain essentially zero unless the transistor enters the active or saturation regions. For example, a neurostimulator powered by a 1.55 V coin cell battery consumes 0.14 ± 0.02 mWh, 2.1 ± 0.3 mWh, and 4.1 ± 1.4 mWh (n = 3, Avg ± SD) when triggered via SWANS, NFC, and BLE, respectively **(Fig. 1E, Supplementary Fig. S1).** While BLE device lifetimes can be improved by decreasing the passive current draw used for advertisement procedures (*29*), this approach effectively limits the feasibility of integrating devices into rapid-acting closed loop systems. In contrast, the SWANS communication structure allows for compact devices with round-the-clock connectivity and effectively no passive current draw.

Each SWANS implant possesses two tissue-interfacing receiving pads connected to the gate pins of a transistor that detect emitted voltage gradient signals in the tissue and facilitate switching. By utilizing different types of transistors switches and passive circuit components, each SWANS implant responds only to a characteristic pulse magnitude, width, and frequency emitted by the wearable hub, enabling the hub to control implants individually. For example, an NPN bipolar junction transistor (BJT) switches on when the potential difference between the base and emitter approaches 0.7 V. Conversely, a PNP BJT turns on around 0.1 V and off near 0.7 V. An N-channel enhancement mode MOSFET transistor turns on at a prescribed threshold voltage, typically from 0.5 V to ≥ 2.0 V. Alternatively, organic electrochemical transistors (OECTs) respond to increasing triggering pulse lengths and rising numbers of sequentially applied pulses by remaining in their on states for proportionally longer periods of time. Additionally, by adding a resistor in between the harvesting pad and the transistor gate of an NPN transistor, it is possible to further increase the required applied voltage gradient necessary for switching. Similarly, by adding a capacitor between the base and emitter of an NPN transistor, it is possible to increase the required pulse length necessary to switch on the implant. With distinct switching criteria for each implanted transistor switch circuit, it is possible to selectively or additively trigger actuation events by applying different voltage gradients to the tissue via wearable source and ground probes.

We engineered the SWANS system to selectively communicate with multiple implanted devices placed anywhere within the body’s conductive tissue by optimizing the voltage magnitude applied to the tissue by a wearable hub as well as the separation distance and surface area of implanted pads used to receive the in-body voltage gradient. In ex vivo studies on chicken breast tissue, the potential difference between receiving pads scaled linearly with the voltage applied by the wearable signal injection probes **(Fig. 1F)**. Applications of 1V, 2V, and 4V pulses yielded received voltage gradients by the implanted pads of 0.20 ± 0.04 V, 0.40 ± 0.06 V, and 0.75 ± 0.07 V (n = 9, Avg ± SD). These pulses provided voltage gradient magnitudes with sufficiently small errors to uniquely regulate the activity of 2N2907 PNP, P55NF06L MOSFET, and 2N2222 NPN transistors, respectively. Increasing the resistance between an NPN base and harvesting pad increased the pulse voltage magnitudes necessary for switching by up to 30% (n=10, **Fig. 1 G**). Adding a capacitor between the NPN base and emitter increased the required pulse lengths necessary for switching from 20 µs to 50-3000µs (n=6-11, **Fig. 1 H**). Combining these two findings, we demonstrated that long pulses with low voltage magnitudes (3.3 V, 2 ms) triggered switches that possessed an added 1 µF capacitor while short pulses with high voltage magnitudes (4 V, 0.05 ms), triggered switches with an added 2.2 kΩ resistor. The pulses that turned on one switch did not turn on the other switch, demonstrating the capacity to uniquely trigger implants (n=8, **Fig. 1 I**). In the case of OECTs, we demonstrated unique triggering patterns in the ex vivo SWANS setup compared to other tested transistors, modulating their switching times based on pulse length and frequency **(Supplementary Fig. S2)**.

The implanted SWANS receiving pads successfully trigger transistor switches when placed across a wide range of orientations, angles, and locations in the tissue. This occurs because the tissue’s conductivity spreads the voltage gradient throughout the entire body. **Fig. 1J** provides a heat map of voltage measurements within an ex vivo chicken breast following an application of a 6V, 1Hz, 200 µs pulse (n=5). The relative orientation of the implanted receiving pads compared to the wearable signal injection probes moderately affects the magnitude of the received voltage gradient. However, this experiment demonstrated the possibility of receiving a > 1 V gradient anywhere between the signal injection and grounding pads, even at angles and locations that are not directly aligned between the two signal transmitter probes.

Ionic communication can trigger millimeter-scale SWANS implants at varying distances and depths away from the wearable signal injection hub. Ex vivo chicken breast studies confirmed that receiving pad separation distances of just 5 mm are sufficient to trigger transistor switches (total communication component size 7 mm x 1 mm), and the required distance between these pads scales with the distance from the signal injection probes (**Supplementary Fig. S3**). Multiple transistors can be uniquely triggered at distances and depths of up to 4 cm away from the signal injection probe (n = 9, **Fig. 1K**); for context, the average skin and subcutaneous fat tissue thickness in a human measures 1.2 ± 0.6 cm (*30*). While further increasing the distance reduces the ability to selectively trigger the implants, all three transistors can be triggered with reduced fidelity at distances and depths of up to 6 cm away from the signal injection probe. Additionally, changing the surface area of the conductive receiving pads between 1-300mm^2^ provided negligible improvement in voltage gradient reception, allowing us to minimize the size of the implanted receiving pads (**Supplementary Fig. S3**). COMSOL simulations provided additional data regarding the effects of pad orientations and separations on electric field generation in both rat- and human-sized tissue models, suggesting that larger distances could be covered by ionic communication in larger models (**Supplementary Fig. S4**). Having determined minimum pad separation lengths, sizes, and orientations for triggering implanted transistors, we next designed wearable and implantable circuits for in vivo SWANS applications.

### A platform for low-power, wireless communication between wearables and implantables

The full in vivo SWANS architecture possesses three key components: (1) a central, wearable hub that coordinates sensor readouts and therapeutic actuation via in-body ionic communication; (2) a microneedle patch that interfaces between the wearable and the body; and (3) an implantable circuit that receives a unique signal and performs a therapeutic action (**Fig. 2A-B, Supplementary Fig. 5**). To enable sensor-guided signaling in vivo, we engineered a wearable flexible printed circuit board (fPCB) containing interchangeable physical or chemical sensors, a control unit, and a configurable voltage output capable of delivering up to 12 V (**Fig. 2C**). These signals are introduced into the tissue via stainless steel microneedle patches, bypassing the highly resistive stratum corneum and directly generating a voltage gradient beneath the skin (**Fig. 2D**). Previous animal and human studies on microneedles and electrical stimulation systems suggest that our employed microneedle lengths and applied current densities will provide a painless, safe, and straightforward user experience (*31–35*). The resulting voltage gradient is then received by a custom-engineered, programmable, implantable fPCB containing two receiving pads, an interchangeable transistor switch, a power source, and an actuator such as a neurostimulator (**Fig. 2E**). In this way, SWANS effectively operates as a low-power, end-to-end communication platform, wirelessly linking wearable sensors to in-body therapeutics.

**Fig. 2.**
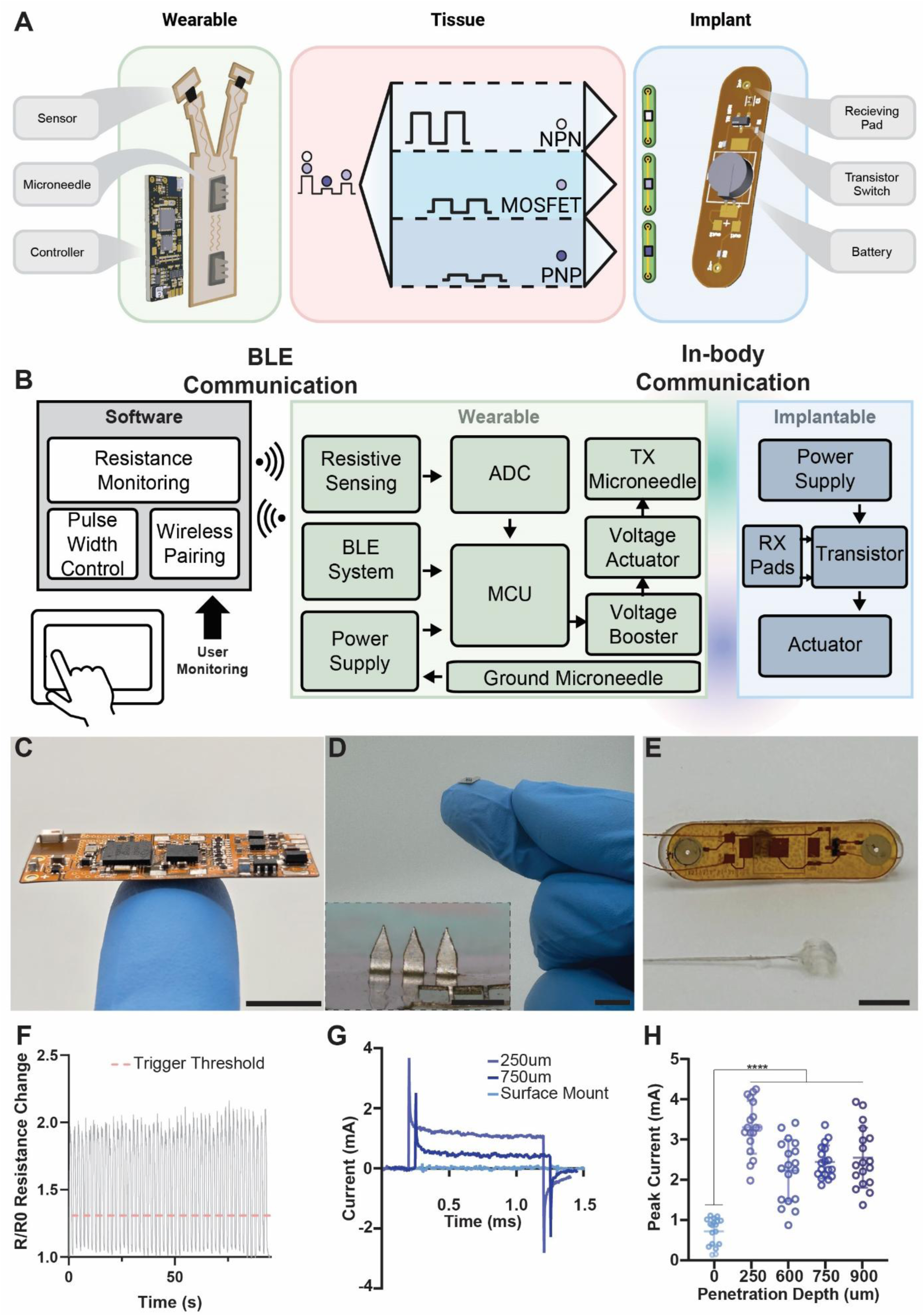
In-body communication architecture. **(A)** A custom flexible printed circuit board (fPCB) continuously monitors a strain sensor and automatically emits a variable (0-12V) square wave pulse to trigger the implantable network. Bioelectronics can be switched individually or synergistically. **(B)** Block diagram for closed-loop control and monitoring of an implantable actuator via in-body ionic communication. **(C)** Photo of wearable fPCB. **(D)** Photo of stainless-steel microneedles that enable the wearable hub’s signals to bypass the stratum corneum (inset scale bar = 1mm). **(E)** Photo of implantable fPCB with voltage receiving pads and a nerve cuff. **(F)** Cyclic testing of a laser-induced graphene (LIG) strain sensor with the threshold for wearable pulsing specified. **(G)** Representative [4 V, 1 Hz, 1 ms] square wave pulses in vivo in rats using skin-interfaced microneedle electrodes. **(H)** Peak injection current across microneedles in vivo in rats (n = 18 across 3 rats, SD plotted). Scale Bars = 1 cm.

The SWANS wearable hub receives inputs from multiple sensors, processes this data, and transmits ionic-communication pulses from 0 V – 12 V into the tissue to control implantable bioelectronics. In our proof-of-concept example, we place two laser-induced graphene (LIG)-based resistive strain sensors (**Supplementary Fig. S6**) on the forelimbs of rats to detect motion intent. The wearable Hub detects when the sensor signal surpasses a given threshold (**Fig. 2F**) and introduces a unique signal into the body depending on the limb that moved. Two implantables with unique transistor switches are placed distally on each hind limb. When an implantable receives its corresponding signal, it stimulates the sciatic nerve on the specified hind leg. Thus, SWANS possesses full control over multiple bioelectronic implants. The implantable circuit’s architecture enables control over a wide array of therapeutically relevant devices in addition to neural interfaces such as pumps and micro-LEDs (**Supplementary Fig. S7**).

The size of the implantable is dictated by the receiving pad separation distance required to trigger transistors at a given depth and the battery requirements for the actuating component. A 2.5 cm pad separation allows for independent control over the three transistors of interest when placed up to 4 cm deep in the tissue; notably, however, the communication components of the device can be scaled down to 7 mm^2^ (7 mm x 1 mm) and still enable switching. This allows for the communication system to fit entirely within a 16 gauge needle and makes it over 5x smaller than the nRF52832 BLE system on chip (6 mm x 6 mm) as well as 1-2 orders of magnitude smaller than many NFC tags (*36, 37*), where working distance scales with coil length.

Due to the highly resistive nature of the stratum corneum, we implemented stainless steel microneedles into the wearable to enable direct electric field generation in the subdermal space, thereby increasing the relative voltage gradient inside the tissue. Microneedles at least 250 µm in height decrease the tissue-electrode interfacial impedance by 4x and increase generated subdermal voltage gradients **(Fig. 2G-H)**.

In vivo ionic communication patterns in rats resembled the trends found during ex vivo chicken breast studies **(Fig. 3A-C)**. Received voltage gradients scaled according to the applied signal and diminished in proportion to the distance between wearable and implanted modules. Application of 7.5 V pulses by a wearable SWANS signal injection probe led to an average received voltage of 0.87 ± 0.16 V, enough to switch an NPN transistor in the subcutaneous space into its active regions during 26/27 trials **(Fig. 3A)**. All PNP and MOSFET transistors actuated in the subcutaneous space at lower signal injection voltages. In the intraperitoneal space, a MOSFET implant entered its active region 9/9 times at an application of 7.5 V, but an NPN transistor only actuated during 6/9 trials **(Fig 3B)**. Receiving pads in the stomach enabled PNP transistor switching at applied voltages of 7.5 V, indicating a possibility for gastrointestinal resident electronics to be regulated via SWANS **(Fig. 3C)**.

**Fig. 3.**
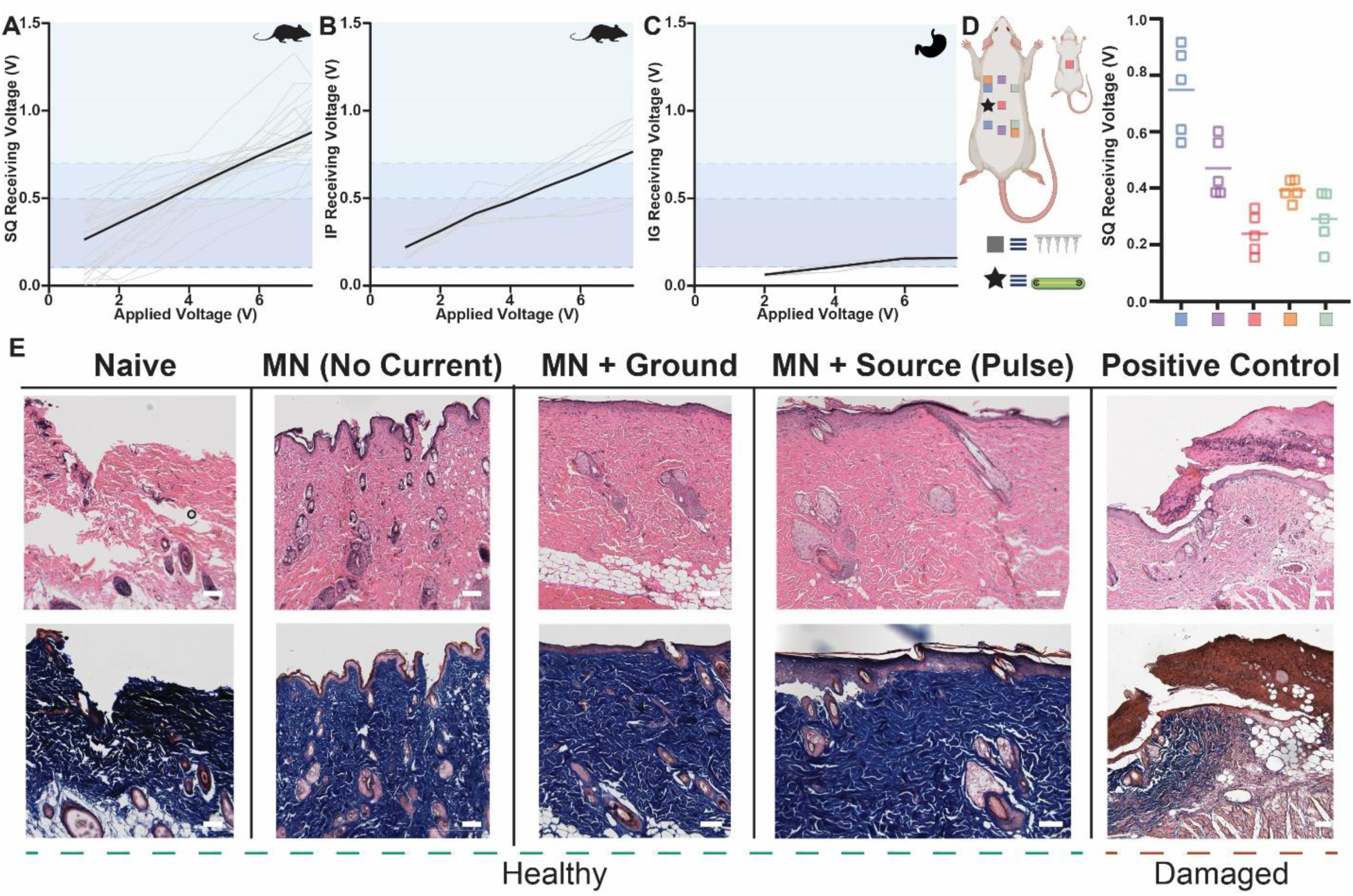
In vivo in body communication generates intrabody voltage gradients without inducing cell trauma. **(A-C)** Voltage measured across a transistor’s gate when applying increasing amplitude pulses across in vivo rat tissue. Receiving pads were placed 2.5 cm apart underneath the skin **(A)** or underneath the peritoneum **(B)** and V_in_ and Ground probes were placed 4 cm apart above the skin. In **(C)**, receiving pads were placed 1 cm apart inside the stomach. Shaded regions denote transistor gate and threshold voltages [black line = avg, grey line = individual tests; (A) n = 27 across 3 rats, (B) n = 9 across 3 rats, (C) n = 10 across 1 rat]. **(D)** Peak received voltage across receiving pads in vivo (n = 5 across 1 rat) when placed at varying orientations to the wearable signal injection probes. Receiving pads were placed 2.5cm apart underneath the skin. V_in_ and Ground probes were placed in spatially varied locations above the skin including in line with the implant (blue), offset from the implant (purple), on the back and belly of the animal (pink), diagonally across the body (orange), and on the opposite side of the implant (green) (n=5; line = avg). **(E)** Representative H&E (top) and Masson’s Trichrome (bottom) stained skin samples upon variable electromechanical treatments performed every other day for one week. Scar tissue formation (brown) is noted when applying 10 V and 100 ms pulses for two straight hours as a positive control, while other histology appears healthy, including when delivering pulses ≤ 10 V, ≤ 10 ms pulses, ≥100 times. Scale Bars = 0.1 mm.

In vivo experiments in rat models demonstrated that SWANS implants successfully actuate when placed at varying orientations, locations, and depths within the tissue. We observed sufficient voltage gradients to trigger a transistor regardless of the spatial orientations of the V_in_ and ground microneedles, including placements horizontally, diagonally, and vertically across the flanks of the rats **(Fig. 3D)**. While the voltage gradient is maximized when TX-RX modules are superposed on each other, as ohmic losses are minimized in this case, current spreads in all directions from the V_in_ source creating voltage gradients in distant areas such as the gastrointestinal tract and the opposite flank of the rat.

Hematoxylin and Eosin as well as Masson’s Trichrome histology staining provided evidence that SWANS 1 Hz pulses of voltages ≤ 10 V for lengths ≤ 10 ms did not generate cellular or tissue damage. We collected rat skin samples after applying microneedles and electrostimulation to the same skin location for at least 100 pulses over the course of one week **(Fig. 3E)**. Previous work on electrical stimulation safety in humans and animal models suggest that the current, pulse length, voltage gradient, charge per phase, and charge density per phase applied by SWANS do not generate skin, neuron, or muscle tissue damage (*32–34*). In our experiment, low current electrostimulation (≤ 5 mA, ≤ 10 V, ≤ 10 ms) induced no visual damage to cells around the microneedle patch and no signs of inflammation. As a positive control comparator, we applied 10 V gradients over 100 ms pulses for 2 hours to generate tissue damage and compare to our test samples.

To demonstrate SWANS’ ability to wirelessly transmit targeted, therapeutically relevant signals across the body, we engineered a bilateral triggerable neurostimulator for rats. SWANS allowed for independent control over multiple implanted devices placed throughout the body in rats. LIG sensors placed on each front paw detected forelimb motion and transmitted this information to a wearable hub which recorded sensor data. When the hub determined that a signal from a given sensor channel exceeded background noise, it activated its voltage booster sending a voltage gradient through the body to trigger the implantable device. **(Fig. 4A**). We confirmed that both NPN and PNP transistor switches on separate implants could be sequentially triggered to deliver sciatic nerve stimulation via our sensor driven wearable hub (n=30 trials over 3 rats **Fig. 4 B-D, F**). In this setup, both NPN and PNP switches triggered leg motion at the higher voltage pulses while only the PNP turned on at the lower pulses. Following the left leg sensor reaching a given threshold, NPN switches turned on with large applied voltage pulses from our wearable device (5+ V), and following the right leg sensor reaching a given threshold, PNP switches turned on with smaller applied voltage pulses (2+ V). Implant activation allowed a sciatic nerve interfacing cuff to receive an influx of power to trigger hind leg motion in the respective limb where the triggered implant was located. As controls, we recorded that the wearable pulse alone without the implant had no impact on hind leg motion at either pulse magnitude, and that the devices alone without an applied pulse had no impact on hind leg motion either (n=30 trials over 3 rats). We captured wearable and implantable electrical data via an oscilloscope, associated sensor data via Bluetooth communication with a nearby tablet, and leg motion data via high speed video. We also created two implants that triggered independently of each other by adding a resistor between the havesting pad and the base of one NPN implant and adding a capacitor between the base and emitter of a second NPN implant (**Fig. 4 E,G** n=30 trials over 3 different rats). Due to tissue impedance variability between rats, these implants and pulses required individualized settings to ensure independent actuation. Only the capacitor (47-68 pF) implant stimuated leg motion during long pulses with medium voltages (1-2 ms, 3.2-8.1 V), and only the resistor implant (422-825Ω) stimulated leg motion during a shorter pulse with higher voltages (0.15-0.25 ms, 4.1-10 V). By forming a bio-integrated network of dispersed wearable and implantable sensors and actuators, the SWANS communication protocol provides a platform technology to enable full-body coordinated control over bioelectronic therapeutic networks.

**Fig. 4.**
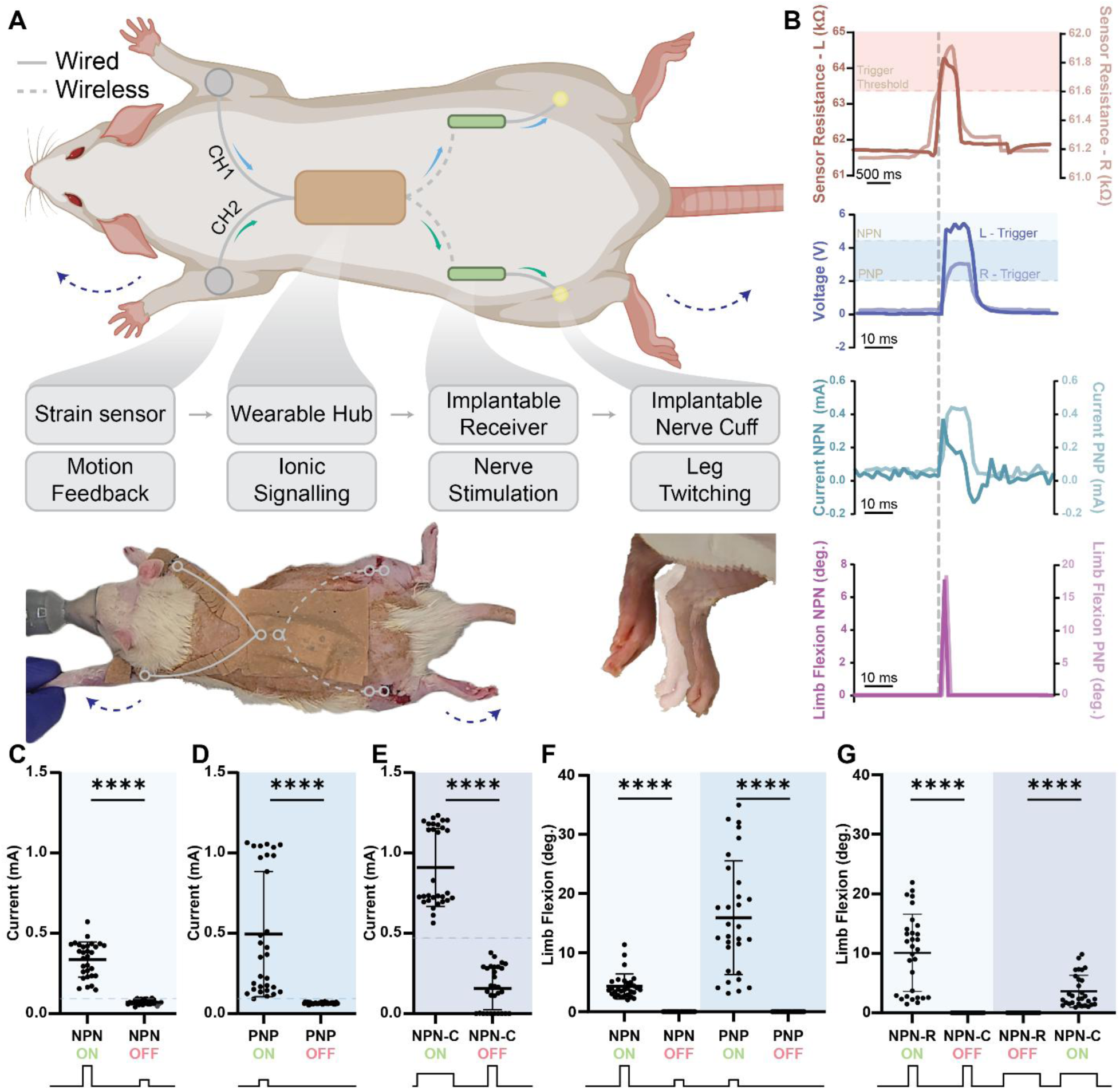
Bio-integrated electronic networks for neurostimulation. **(A)** Schematic and image of rat outfitted with SWANS neurostimulator system with representative image of leg twitch from resting [translucent] to flexed [opaque]. **(B)** Upon stimulus detection on either forelimb, the wearable board outputs different pulse magnitudes and widths, which trigger different implants to actuate. **(C)** Current outputs from NPN implants using high V short pulses, **(D)** PNP implants using low V short pulses, and **(E)** NPN + Capacitor (NPN-C) implants using longer medium V pulses. Points above dotted lines represent presence of leg flexion. **(F)** Triggering the NPN and PNP implant combination led to sequential triggering of PNP only followed by NPN + PNP with increasing voltage when placed in the same animal. **(G)** NPN + Resistor (NPN-R) implants only triggered with High V short pulses and NPN-C implants only triggered with medium V long pulses when placed in the same animal. Pulse lengths and magnitudes varied across animals due to variance in skin thickness and tissue impedance, as described in Materials and Methods. (n=30 over 3 rats with different rats tested in each panel, Avg and SD plotted, **** indicates p<0.0001 in unpaired t-test with Welch’s correction).

## Conclusion

SWANS reads out wearable sensor data and wirelessly triggers networks of implantable therapeutic devices via ionic conduction. Here, we demonstrate sensor-mediated neuromodulation in rats, and our data suggests that future SWANS iterations could potentially restore limb locomotion (*38, 39*), actuate smart pills (*40, 41*), trigger drug delivery reservoirs (*16, 17, 42*), switch on optogenetic probes (*43, 44*), and modulate living cellular implants (*45–47*) as well as control other bioelectronics throughout the epidermal, subcutaneous, intraperitoneal and gastrointestinal spaces. SWANS implantables require an order of magnitude less power than BLE and NFC without necessitating an external antenna. SWANS excels in applications where infrequent triggering and low data transmission rates are required, as implants draw near-zero power consumption but have limited data transfer rates. The communication components alone can be scaled down to fit inside of a 16 gauge needle, are over 5x smaller than the nRF52832 BLE system on chip, and are orders of magnitude smaller than many NFC coils used in vivo (*36, 37*), opening a new sandbox for biomedical engineers. In the future, it may be possible to increase the complexity of data transmission to more than on/off signaling by adding a microcontroller that reads out pulsed patterns, but this will come at the cost of reduced battery life and increased size. While preliminary histology studies suggest that SWANS current injections do not generate tissue damage, long-term pain measurement and tissue health studies in humans are required to determine the safety of the SWANS system. Future studies in large animal models will quantify generated voltage gradients over larger decimeter-scale distances, with previous studies in humans and large animals suggesting the scalability of ionic communication (*23, 26*). Additionally, utilizing conductive gels on the skin may reduce the impedance across the stratum corneum, potentially eliminating the need for microneedles in cases that require lower voltage gradients (*48*). The communication efficacy achieved with this platform technology suggests that SWANS could support a new generation of dispersed and coordinated bioelectronic therapies, opening the possibility to integrate and network the countless numbers of wearable sensors and therapeutic devices being engineered for personalized medical treatments.

## Acknowledgments

We are grateful to all members of the Abramson Lab for their helpful discussions. We thank Nathaniel Kirkpatrick, Richard Noel, and Laura O’Farrell for their help with animal experiment design. We are thankful to Alana Mermin-Bunnell for early discussions on the project. We thank the Parker H. Petit Institute for Bioengineering and Bioscience, School of Chemistry and Biochemistry, and the Office of the Executive Vice President for Research for their generous support. This work was performed in part at the Georgia Tech Institute for Matter and Systems, a member of the National Nanotechnology Coordinated Infrastructure (NNCI), which is supported by the National Science Foundation (Grant ECCS-2025462).

## Funding

Georgia Tech Institute for Electronics and Nanotechnology 1000x seed grant (A.A., W.- H.Y.)

NSF CAREER Award #2439870 (A.A.)

NSF GRFP fellowship (R.G, J.J., S.H.)

Georgia Tech President’s Undergraduate Research Award (A.K., A.C.)

Georgia Tech Petit Scholars Research Award (E.F., A.C.)

Beckman Coulter Scholarship (A.C.)

NIH MIRA grant R35GM150689 (A.A.)

WISH Center grant from the Institute for Matter and Systems at Georgia Tech (W.-H.Y.)

NSF NRT-FW-HTF grant #2345860 (J.D., W.-H.Y.)

Research Foundation of Korea (A.A.).

## Author contributions

R.G. and A.A. created the idea and designed the experiments supporting the development of SWANS. R.G., E.F., J.J., D.A., A.K., A.C., S.H., and J.D. performed the ex vivo characterization studies on ionic tissue communication in chicken breast tissue. C.C., A.G., and R.G. fabricated and characterized organic electrochemical transistors. R.G., E.K., Y.J.L., and G.B. designed and fabricated the printed circuit boards. R.G., E.F., J.P., and J.J. performed or assisted with in vivo experiments characterizing ionic tissue communication in rats. R.G., Y.J.L., G.B., and A.A. performed or assisted with in vivo neuromodulation experiments. A.A. and W.-H.Y. provided funding and supervised the project. All authors contributed to writing and editing the manuscript.

Conceptualization: RG, AA

Methodology: RG, AA

Investigation: RG, YJL, GB, EK, EF, JJ, DA, AK, AC, SH, JD, CC, JP

Visualization: RG, AA, JJ, GB, YJL, CC, AG

Funding acquisition: AA, WHY

Project administration: RG, YJL, AA, WHY

Supervision: AA, WHY, AG

Writing – original draft: RG, EF, JP, JJ, DA, AK, AC, SH, JD, CC, GB, YJL, EK, AA, WHY, AG

Writing – review & editing: RG, EF, JP, JJ, DA, AK, AC, SH, JD, CC, GB, YJL, EK, AA, WHY, AG

## Diversity, equity, ethics, and inclusion [optional]

We support inclusive, diverse, and equitable conduct of research.

## Competing interests

A.A., R.G., Y.J.L., and W.-H.Y. are inventors on provisional patents describing the technology presented in this manuscript. A.A.’s full list of competing interests can be found at site.google.com/view/alex-abramson-coi.

## Data and materials availability

All data are available in the main text or the supplementary materials.

## Notes

### Competing Interest Statement

A.A., R.G., Y.J.L., and W.-H.Y. are inventors on provisional patents describing the technology presented in this manuscript. For A.A., a full list of competing interests can be found at site.google.com/view/alex-abramson-coi.

### Summary of Updates

Updated text and figures for clarity

