## Supplementary for "An artificial nervous system for communication between wearable and implantable therapeutics"

**The PDF file includes:**

Materials and Methods
Figs. S1 to S7

### Materials and Methods

Transistor switching experiments with ex vivo chicken breast

Chicken breast (150-200g, Heritage Farms) was soaked in PBS (VWR) for up to two hours. Using a function generator (Siglent - SDG2000X), 1 Hz monophasic pulses of various amplitudes were applied across 13 cm of tissue. Voltage generation and harvesting were accomplished with hypodermic needles (BD) or laser-cut stainless-steel shim (McMaster-Carr) as described in "Fabrication of Microneedles" below. Receiver pads were positioned on the tissue surface at various separations between the signal injecting and grounding probes including pad separations of 9, 5, 3, and 0.5cm.

Applied and harvested voltages, along with current through a 100Ω shunting resistor, were measured using an oscilloscope (Rigol-MSO5000). Current was measured by probing both sides of the resistor, smoothing the data to remove noise by taking a 10-point moving average, and measuring the peak difference between the voltage measurements during the pulse. The max current was reported ex vivo; for in vivo experiments, the max average current over a 50 μs period, the time required to enable a leg twitch, was recorded. 3 transistors including a 2N2222 (NPN), 2N2907 (PNP), and a P55NF06L (MOSFET) were connected in series with a 3V battery and an LED to determine required applied voltages to reach the active and saturation regions of each transistor. To construct a visual representation of the generated voltage gradients, potentials were collected at various locations across the tissue with respect to a grounding tissue-interfaced electrode.

Surface area experiments with ex vivo chicken breast

Chicken breast was soaked in PBS (VWR) for up to two hours. Using a function generator, 1 Hz monophasic pulses of various amplitudes were applied across 13 cm of tissue. 10 $mm^2$ stainless steel electrodes were directly connected to the function generator. Various sized electrodes (1, 2, 3, 5, 7.5, and 10 $mm^2$) were placed 9cm apart to capture a portion of the generated gradient. Harvested potentials were recorded with an oscilloscope.

Device Fabrication

**Fabrication and characterization of the Wearable package:** A wearable pulse generation system was developed, featuring a flexible PCB (fPCB) circuit layered with polyimide-copper for skin conformity and flexibility. The wearable device connects with an Android application, allowing contactless control of pulse delivery and custom adjustments of parameters such as frequency (0.01-1 Hz) and pulse width (1-1000 ms). The fPCB utilizes the nRF52832 microcontroller (MCU) architecture, selected for its industry standard low-power Bluetooth capabilities. This wireless system integration allows user-defined control via the mobile app for open/close/remote controls.

The device's voltage range spans from 1V to 12V, powered by a custom-configured, controllable DC-DC step-up voltage regulator and booster. A DC-DC boost converter that steps up the LiPo battery voltage (3.7V nominal) to the custom stimulation voltage (0-12V adjustable). The design is customizable with stimulation scenarios using both transistors and MOSFETs, regardless of NPN or PNP types, allowing for programmable voltage amplitude, pulse width, and frequency to generate dynamic pulses. A set of power switch drivers generates user-programmable monophasic or biphasic pulses that can be controlled by firmware code, making the device fit for various stimulation protocols. The system-on-chip placed at the center of the PCB processes executing the main control algorithm, reading sensor data, and driving the stimulation pulses.

To facilitate input-triggered reading/actuation, an ADS1292 24-bit analog-to-digital converter (ADC) is integrated, interfacing with resistive sensing elements. These elements detect mechanical stimulation, such as strain changes, and can initiate pulse generation in response to physical cues and for biofeedback system. The system is designed with minimal traces length and physical hardware for data communication, operation, and critical signals. The ground planes are separated to reduce electromagnetic interference and/or logic interference and/or control interference. Flexible serpentine traces interface with the LIG sensor and are designed to maintain mechanical compliance and reduce stress at joints of the pads. The strain that was continuously measured by a Wheatstone-type resistor network of resistor branches with adjustable reference thereby allows differential resistance sensing and fine-tuning of offset and gain for further calibration, calibration curve, and measurement precision.

All firmware and software development were conducted in a Windows desktop environment. Firmware is compatible with Nordic nRF52 platform Soft Device stack for BLE functionality and acknowledgement. The companion mobile application was implemented in Kotlin (2022) for Android devices, tested performed on a Samsung Galaxy S8 tablet.

**Fabrication and characterization of the Implantable package:** The implantable device, measuring 25 by 10 mm, is designed to function as a subdermal signal receiver and programmable output stimulator. The device is powered by a 6.8 mm diameter 1.55V silver oxide coin cell battery (Duracell 377), a 4.8 mm diameter 3V lithium coin cell battery (Seiko Instruments, ML414H IV01E), or a 10 mm 3V lithium coin cell battery (Fuspower, CR1025) selected for their compatibility with gate voltage requirements and operational lifetime. The entire implantable fPCB is encapsulated in PDMS (Sylgard 184, Dow) using a 2-piece mold that ensures that gold contact pads for signal harvesting interface with surrounding tissue. Loctite 4311 medical grade epoxy coating could further isolate the electronics from the surrounding tissue. The stimulation is programmable through the GPIO of the wearable's microcontroller, allowing dynamic control over the applied stimulation parameters. For signal modulation, the implantable system supports selective use of NPN, PNP, or MOSFET components to regulate the gate voltage and control the current passing through the implantable. It also possesses location to add resistors between the harvesting pads and transistor gate, as well as capacitors between the base and emitter as needed.

**Fabrication of Microneedles***:* 75μm stainless-steel shim (McMaster-Carr; 316SS) was laser-cut using a femtosecond laser-cutter (60W, 75% power, 150 repetitions, or 4W, 60 kHz, 100% power, 45 repetitions, Optec). Microneedles were manually prodded out using a razor blade or hypodermic needle. Microneedles were cleaned and electropolished using a 6:3:1 glycerin:85% phosphoric acid: DI water.

**Laser-Induced Graphene sensor fabrication and characterization.** The fabrication of the uniaxial LIG sensor began with the synthesis of styrene-ethylene-butylene-styrene (SEBS) in a dissolution concentration at 5%, homogenized on a hotplate at a determined temperature for a precise duration to ensure uniform solution characteristics. The stainless steel (SS) substrate was prepared by spraying it with an anti-stick aerosol (Ease Release 200, Reynolds Advanced Materials) to ease subsequent material removal. SEBS solution was then drop-cast onto the SS substrate and cured at 35°C for 2 hours, resulting in a uniform SEBS film. Laser micromachining using the OPTEC Femtosecond Laser was employed to achieve precise film patterning, after which the negative material was carefully removed. For the LIG component, a polyimide (PI) sheet of 75 um thickness underwent $CO_2$ laser cutting at 5% power (Trotec Speedy laser, 5% power, 0.2 speed, 16 kHz), which induced the formation of graphene layer on its surface. The LIG strip was transferred onto the stretchable SEBS substrate and PI/Cu interconnect layer through mechanical stamping, with pressure applied at 100N over ten repetitions to ensure strong adhesion. The interconnect and LIG components were then sealed together by SEBS-SEBS film adhesion, which relies on Van der Waals forces for a secure bond, achieving both durability and stretchability without increasing overall thickness. Finally, the exposed copper pads of the PI/Cu interconnect were soldered to the 2CH lead from the device, and the entire assembly was bonded to a fabric adhesive for secure mounting onto biological substrates.

To measure the resistance during stretching, we attached the samples to a homemade stretching station and connected the samples to a tabletop multimeter. A custom LabView program simultaneously collected force, displacement, and resistance data. Samples were stretched at 1mm/s up to 20% strain for up to 50 cycles.

To characterize on-body performance, a laser-induced graphene (LIG) sensor is affixed to the skin of the small animal using brown medical tape. The LIG strain sensor displays a gauge factor of 5-10 demonstrates large relative resistance changes with only small mechanical deformations (<5% strain). In practical terms, these sensors can detect subtle limb or finger movements down to sub-millimeter displacements. The minimum resolvable strain can be as low as 0.5% in the presence of environmental noise.

**Fabrication of Bi-layer Copper Flexible 2CH Interconnect:** Sylgard 184 (Dow) with a 10:1 PDMS to crosslinker ratio was prepared. The prepared PDMS was subsequently spin-coated onto a stainless-steel substrate to ensure a smooth surface, set at a speed of 1000 rpm for a duration of 35 seconds, with a ramp period of 5 seconds included. The elastomer was then cured at 120°C on a hotplate for 30 minutes. A copper (Cu) foil was laminated onto the PDMS surface; due to the inherent Van der Waals forces, no additional adhesive was required to achieve adhesion, attributed to the high adhesion energy of PDMS molecules that promote stable interfacial contact.

Following the initial lamination, a layer of low-modulus silicone elastomer with high surface energy (Ecoflex-GEL, Smooth-On Inc.) was applied on the laminated Cu film via spin coating at 1500 rpm for 35 seconds. The applied elastomer was then cured at 65°C for 1 hour, providing a uniform insulating layer. Subsequently, a second Cu foil was laminated onto the cured elastomer layer, resulting in a bi-layer configuration of parallel Cu films separated by a thin elastomer layer that serves as an insulator to prevent electrical shorting.

To pattern the Cu-elastomer-Cu composite, laser micromachining was performed using an OPTEC Femtosecond Laser system. A multi-step cutting process was employed to minimize substrate damage when processing the relatively thick composite layer. Using the top Cu film as the baseline focal point, patterning was conducted at 65% laser power for five repetitions. The focal point was then lowered by 250 μm, and the cutting process was repeated. This sequence was iterated three times to ensure full penetration through the composite to the bottom Cu layer. After laser processing, negative material was carefully removed, and the resulting bi-layer interconnect was laminated with PVA water-soluble tape (5414 Water-Soluble Wave Solder Tape, 3M) to facilitate its transfer from the PDMS surface for further integration onto fabric adhesive substrates.

**Fabrication of PI/Cu Flexible Interconnect:** The PDMS precursor was prepared with Sylgard 184 (Dow) at a 10:1 PDMS to crosslinker ratio and then spin-coated onto a stainless-steel substrate at 1000 rpm for 35 seconds, including a 5-second ramp, to achieve a smooth elastomer layer. Curing was conducted on a hotplate at 120°C for 30 minutes. After curing, Cu foil was laminated onto the PDMS surface, where it adhered naturally due to Van der Waals forces, facilitated by the high adhesion energy of PDMS molecules.

Following Cu lamination, a heat-activated polyimide (PI) adhesive film (Pyralux LF, DuPont) was applied onto the Cu foil, and bonding between the Cu and PI layer was achieved by heat pressing at 400°C for 15 seconds. The thermoplastic adhesive was activated under heat, forming a cross-linked polymer matrix between the PI and Cu layers, thereby creating a permanent bond.

The PI/Cu composite film was subsequently patterned using laser micromachining with an OPTEC Femtosecond Laser at 75% power, applied in five repetitions. The laser cutting parameters were meticulously optimized to remove material effectively while preventing thermal damage, which could introduce carbonized impurities along the patterned edges. Upon completion of the laser patterning, the negative material was removed, and the remaining bi-layer interconnect was laminated with PVA water-soluble tape (5414 Water-Soluble Wave Solder Tape, 3M) to facilitate detachment from the PDMS substrate and subsequent transfer to a fabric adhesive for final mounting.

Voltage Receiving Simulation

$\nabla \cdot D = \rho_v$ (1)

$E = -\nabla V$ (2)

Geometry generation and E-field distribution analysis simulations were performed using the commercial finite element method (FEM) software, COMSOL Multiphysics version 6.2. (Burlington, MA, USA) To simplify the modeling procedure, a pair of microneedle pads, an implant-receiving pad, and the skin were implemented in the simulation. The relative permittivity of the skin was 6000, and the electrical conductivity was set as 0.1 S/m. The microneedle pads were selected as iron, which had a relative permittivity of 4 and the electric conductivity of $10.2 \times 10^6$S/m referenced from the COMSOL material library. The receiving pad from the implantable device was set to gold, which has a relative permittivity of 5, and electrical conductivity was set as $45.6 \times 10^6$S/m, also referenced from the COMSOL material library. The electrostatic equation was used for the electric potential simulation. It is defined by Eq. (1) and Eq. (2).

OECT Fabrication

OECT devices with dimensions of L = 5 μm and W = 400 μm were fabricated in the cleanroom environment. Source and drain electrodes were defined using photolithography with AZnLOF2020 photoresist and a mask-less aligner (Heidelberg MLA 150). A 10 nm layer of Ti followed by a 100 nm layer of Au was deposited by using an e-beam evaporator (Temescal FC2000). Subsequently, a first layer of parylene-C was deposited using a Specialty Coating Systems PDS 2010, employing Silane A174 (Sigma-Aldrich) as an adhesion promoter. A sacrificial layer of 2% MICRO-90 soap (VWR) in DI water was spin-coated onto the wafer before depositing a second layer of parylene-C. The channel and electrode pads were then defined using the same maskless aligner with AZ10XT photoresist and a reactive ion etching process (SAMCO 230iP). Channels were formed by spin-coating a 5 mg/mL solution of a semiconducting polymer, poly[(2,6-bis(thiophen-2-yl)-3,7-bis(9-octylnonadecyl)thieno[3,2-b]thieno[2',3':4,5] thieno[2,3-d]thiophene-5,5'-diyl)-alt-(3,6-bis(thiophen-2-yl)-2,5-bis(8-octyloctadecyl)pyrrolo[3,4-c]pyrrole-1,4(2H,5H)-dione)-5,5'-diyl] or DPP-4T (Corning Incorporated), onto the devices at 2000 rpm for 35 s via dynamic spin-coating. Following a peel-off step, the devices were baked inside a glovebox at 120 °C for 30 min and allowed to cool for at least 1 h before characterization. A PDMS sheet was cut to form a well and used to contain the 1-ethyl-3-methylimidazolium bis(trifluoromethylsulfonyl)imide (Sigma-Aldrich) electrolyte. Transfer characteristics, as well as responses to a pulse burst, of the OECT devices were then measured with an Ag/AgCl pellet as the gate using a Keithley 4200A. Data were plotted using OriginLab software.

In vivo signal injection characterization

All animal procedures were approved by the Georgia Institute of Technology Institutional Animal Care and Use Committee and conducted in accordance with Georgia Institute of Technology animal facility guidelines. Adult male and female Sprague Dawley rats (200-300g; Charles River

Laboratories, Wilmington, USA) were maintained on a 12h:12h light:dark cycle (lights on at 7:00). Subjects were anesthetized and placed in a lateral recumbent position. 2 patches of skin separated by 4cm were shaven and cleaned aseptically. Various electrodes (surface mounted – 0µm penetration, 300µm, 600µm, 750µm, 900µm, and 1600 µm) were connected to a function generator in series with a 100Ω resistor. 1 Hz, monophasic, 4V pulses were used to generate voltage gradients across the rat and the injected current was measured.
Additionally, the injected current using 900µm microneedles was quantified for a range of applied voltages from 0.5V – 10V.
**Transistor switching experiments in vivo:** Subjects were shaven and prepared for aseptic surgery using 3 cycles of chlorohexidine and 70% ethanol application. A 1cm incision was made perpendicular to the animal torso halfway between the animal's fore- and hindlegs. 1mm diameter gold coated PDMS pads (300rpm, 10:1 ratio cured at 70°C; SYLGARD 184, Dow, Midland, USA) were placed inside the subcutaneous space on either side of the incision at a separation of 2.5 cm. The incision was closed temporarily with sutures (4-0 silk, Covidien) to prevent the subcutaneous space from drying. Microneedles were applied onto the skin 0.75 cm away from each of the implanted conductive pads and connected directly to a function generator. Applied and harvested voltages were measured as described previously.
Additionally, transistor switching was performed by placing these gold pads on the peritoneal lining simulating an implantable device placed in the intraperitoneal space. Finally, the gold pads were placed inside the stomach by exteriorizing the stomach, making a 0.5cm incision and placing the pads inside the stomach. Afterwards, the stomach was placed back inside the intraperitoneal space.
Applied and harvested voltages, along with current through a 100Ω shunting resistor connected in line with the nerve cuff, were measured using an oscilloscope. The average maximum current passing through the resistor and nerve cuff over 30 µs during each trial was recorded.
NPN-R and NPN-C transistors that demonstrated independent switching in vivo required different pulse lengths, resistances, capacitances, voltage magnitudes, microneedle lengths, and microneedle locations depending on the animal. In each animal, these values remained the same throughout the entire data collection experiment to ensure that the switching only occurred due to the changes between two defined pulses. In animal 1, a female rat with thinner skin, the settings were as follows: 422Ω Resistor; 68 pF Capacitor; 10 V / 250 µs pulse to turn on the NPN-R implant; 6.5 V / 2 ms pulse to turn on the NPN-C implant; 900 µm microneedle length. In animal 2, a male rat with thicker skin, the settings were as follows: 825Ω Resistor; 47 pF Capacitor; 10 V / 100 µs pulse to turn on the NPN-R implant; 8.0 V / 1.2 ms pulse to turn on the NPN-C implant; 1600 µm microneedle length. In animal 3, a male rat with thicker skin, the settings were as follows: 825Ω Resistor; 47 pF Capacitor; 4.1 V / 150 µs pulse to turn on the NPN-R implant; 3.2 V / 2 ms pulse to turn on the NPN-C implant; 1600 µm microneedle length.
**Closed-loop demonstration via in-body communication**: Subjects were shaven and prepared for aseptic surgery using 3 cycles of chlorohexidine and 70% ethanol application. Female Sprague Dawley rats (200-300g, Charles River) were anesthetized with 2% isoflurane in balanced oxygen. Under sterile conditions with external body warming, a 2-cm incision was made on the left and right thigh. About 1 cm of the sciatic nerve proximal to the tibial and peroneal bifurcation was inserted into the nerve cuff electrode (NC-2-2-125SS-1.5-2-00-300-00, MicroProbes). Nerve cuff leads were tunneled through the muscle layer (closed with 4-0 silk sutures (Ethicon)) into the subcutaneous space where the implantable FPCB was placed. The skin was secured with sutures.

The subject was placed in a prone position and was outfitted with the wearable FPCB including microneedles and a soft strain sensor. A soft tether was applied to each of the forelimbs and the tether was controlled by the researcher to simulate extension of the forelimb and a natural walking motion. Electrical characterization of the wearable and implantable FPCBs was completed using an oscilloscope. Resistance measurements were collected using a tabletop multimeter (Keithley). Frames from video recordings set up for monitoring each of the hindlegs were extracted and used to determine the degree of limb flexion.

The wearable and implantable devices operate in a closed-loop configuration for adaptive therapeutic response. The wearable device continuously monitors motion through a Laser-Induced Graphene (LIG) sensor, capturing motion range and detecting boundary conditions. Upon detecting a boundary condition change, the wearable's pulse generator activates a voltage boost, sending an electrical trigger to the implantable device.

This closed-loop control enables targeted neurostimulation via a microneedle interface in the implantable system, precisely adjusting the stimulation intensity in response to user movement or external stimuli. The implantable device, acting as an electrical gate, modulates the stimulus through programmable parameters, ensuring effective and responsive therapy delivery within the in vivo model.

<u>Tissue Health and Histological Examination Studies of applied voltage</u>

For examination studies, microneedles were applied to rat skin in vivo and stimulated with 100 pulses by the wearable FPCB or a function generator and imaged with an optical microscope immediately after the pulses and again after 3 hours. For hematoxylin/eosin (H&E) staining and Masson's Trichrome staining, the tissue was either paraffin embedded or frozen. For paraffin embedded samples, tissue samples were fixed overnight in 10% formalin buffer, then dehydrated by an automatic tissue dehydration system. The dehydrated tissue was embedded in paraffin and sectioned at 5um thickness by rotary microtome. For frozen samples, rat skin tissue samples were immediately embedded in optimal cutting temperature compound (Sakura Finetek, CA), frozen with dry ice, and sectioned at 5 μm thickness with a cryostat. Stained slides were imaged using an inverted microscope.

**SI 1. BLE current consumption profiling and power consumption comparison across communication methods.** (A) Representative BLE current profile measured using the nRF Profiler, illustrating average and peak current consumption during discrete operational stages: pre-processing, ramp-up, standby, radio startup, RX (receiving), radio switching, TX transmission, and post-processing. (B) Comparative summary of theoretical and experimental power consumption metrics (mean ± SD, n = 3) for Implantable transistor-based system, Bluetooth Low Energy (BLE), and Near Field Communication (NFC) communication. The table highlights standby currents, active currents, daily power consumption, and estimated battery lifetimes for each technology, demonstrating the substantial energy-saving advantages of the implantable transistor-based system compared to BLE and NFC.

**A**

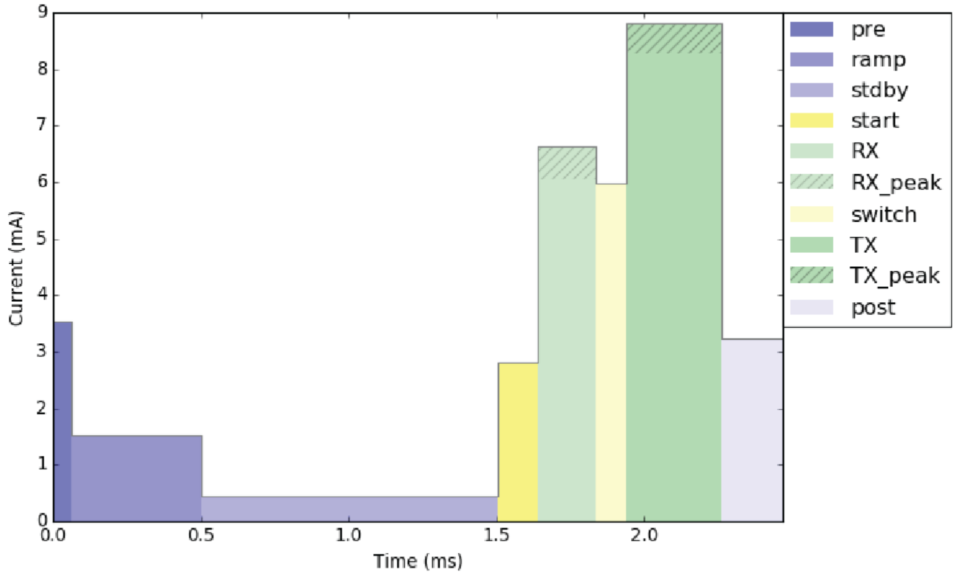


| Stage | Description | Time (ms) | Length (us) | Avg. current (mA) | Peak current (mA) |
|---|---|---|---|---|---|
| pre | Pre-processing | 0.0 | 61 | 3.5 | |
| ramp | Standby + HFXO ramp | 0.1 | 440 | 1.5 | |
| stdby | Standby | 0.5 | 1004 | 0.4 | |
| start | Radio startup + CPU | 1.5 | 133 | 2.8 | |
| RX | Radio RX | 1.6 | 200 | 6.0 | 6.6 |
| switch | Radio switch | 1.8 | 102 | 6.0 | |
| TX | Radio TX | 1.9 | 323 | 8.3 | 8.8 |
| post | Post-processing | 2.3 | 205 | 3.2 | |
| | System On IDLE | 2.5 | 1997.5 ms | 2.0 uA | |
| **Total** | | | **2000.0 ms** | **5.4 uA** | |

**B**

| Scenario | Parameter | Theoretical Consumption | Experimental Consumption (Mean ± SD, n=3) | Estimated Battery Life (Mean ± SD) |
|---|---|---|---|---|
| **In Body Communication (IBC)** | Standby Current | 1 μA × 1.55 V = 1.55 μW | 1.2 ± 0.1 μW | 281 ± 10 days |
| | Active Current | 5 mA × 1.55 V = 7.75 mW | 7.8 ± 0.5 mW | |
| | Daily Consumption | 0.134 mWh | 0.140 ± 0.02 mWh | |
| **Bluetooth Low Energy (BLE)** | Standby Current | 2 μA × 3.0 V = 6 μW | 6.5 ± 0.5 μW | 40 ± 5 days |
| | Active Current (TX) | 8.3 mA × 3.0 V = 24.9 mW | 25 ± 1 mW | |
| | Daily Consumption | ~1.5 mWh | 1.5 ± 0.2 mWh | |
| **Near Field Communication (NFC)** | Standby Current | 5 μA × 3.3 V = 16.5 μW | 17 ± 1 μW | 31 ± 2 days |
| | Active Current | 20 mA × 3.3 V = 66 mW | 65 ± 2 mW | |
| | Daily Consumption | 2 mWh | 2.1 ± 0.3 mWh | |

**SI 2. Organic electrochemical transistors (OECTs) enable digital switching and long-term drain current after pulsing.** (A) OECT response to 10 pulses of 100 ms pulses [0, -0.8 V, 50% duty cycle (ED)], $V_{ds}$ -0.6 V. (B) OECT response to 10 pulses of 1 sec [0, -0.8 V, 50% ED], $V_{ds}$ = -0.6 V (C) First 10 transfer curves of a DPP-4T-based OECT device in EMIM-TFSI. (D) OECT response to 1 pulse of 100 ms pulse [0, -0.8 V, 0.5 Hz] applied to ex vivo chicken breast model, $V_{ds}$ -0.6 V. (F) OECT used to switch on/off an LED using an ex vivo chicken breast model.

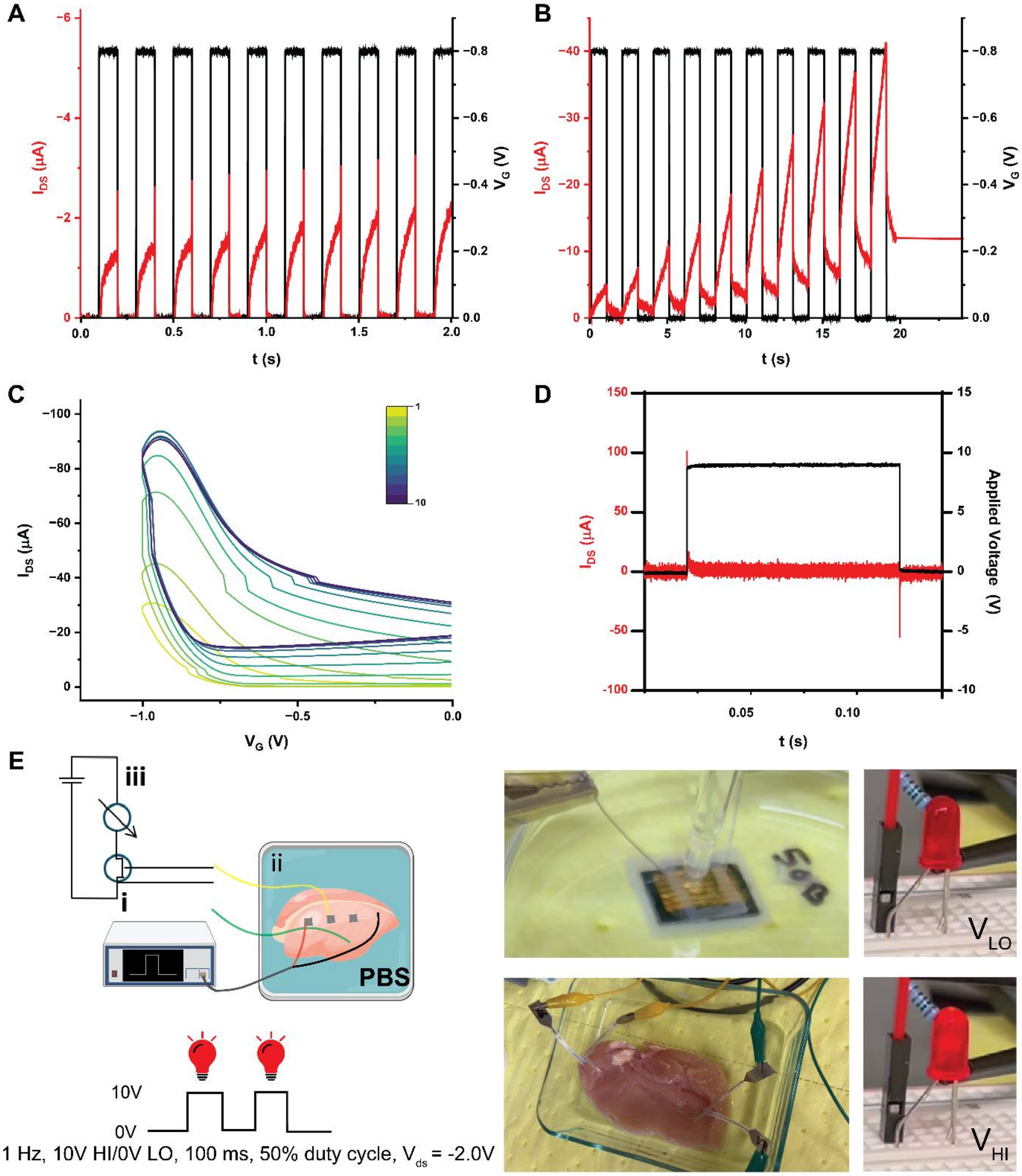

**SI 3. Continued ex vivo characterization of in-body communication and transistor switching.** (A) Receiving pads are maintained at a constant distance of 3 cm, while the signal injection pad are varied from 1, 2, and 5 cm from the receiving pad. (Black lines = mean, n=3). (B) Various receiving pad surface areas at a constant applied voltage of 7 V and separation of 3 cm between receiving pads.

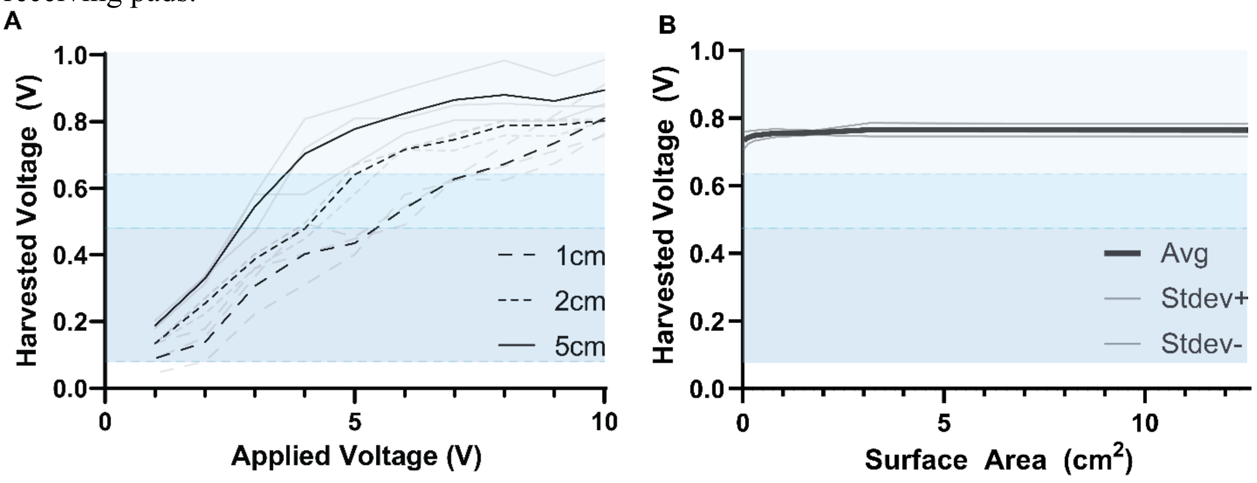

**SI 4**. **COMSOL simulations of electric-field generation in rat- and human-model tissue.** (A) Transmitter pads create an electric field and generate voltage gradients across centimeter-scale model tissue. Receiving pads detect these voltage gradients as a function of pad orientation, spatial position, and separation. (B) Electric field generation in rat-model tissue. (C) Electric field generation in human-model tissue. (Next page) Voltage gradients as a function of pad orientation, spatial position, and separation at applied voltages of 3 V and 7 V. Scale bar = 1 cm.

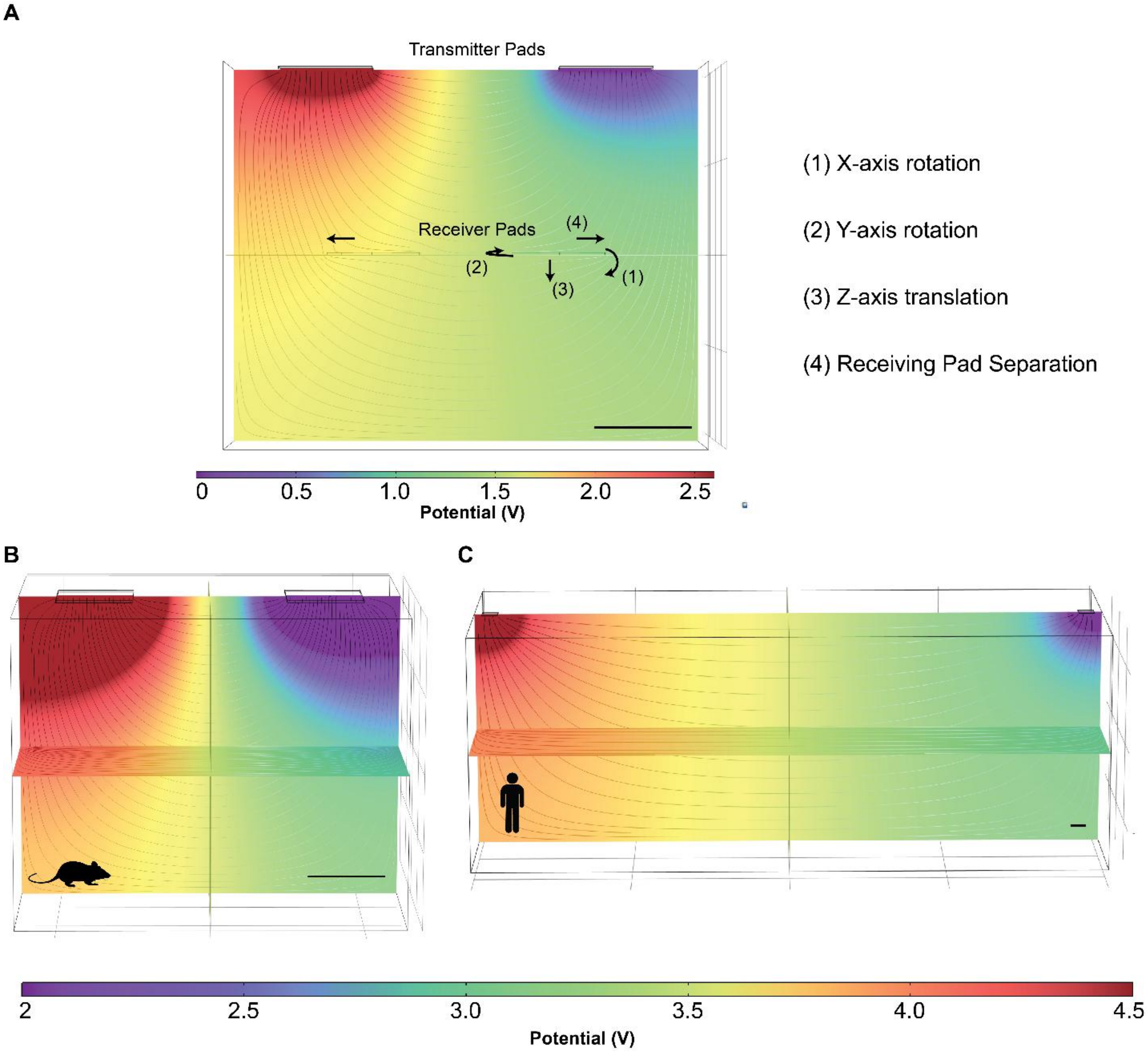

| **Receiving Pad Transformation** | **Applied Voltage (V)** | **Configuration** | **Receiving Voltage (dV, V)** |
|---|---|---|---|
| (1) X-axis Rotation | **3** | **In-line (0°)** | **0.5** |
| | | Offset (30°) | 0.5 |
| | | Offset (60°) | 0.49 |
| | 7 | In-line (0°) | 1.18 |
| | | Offset (30°) | 1.17 |
| | | Offset (60°) | 1.15 |
| (2) Y-axis Rotation | **3** | **In-line (0°)** | **0.50** |
| | | Offset (30°) | 0.44 |
| | | Offset (60°) | 0.26 |
| | 7 | In-line (0°) | 0.50 |
| | | Offset (30°) | 1.02 |
| | | Offset (60°) | 0.60 |
| (3) Z-axis Translation | **3** | **2 cm** | **0.50** |
| | | 3 cm | 0.30 |
| | | 4 cm | 0.24 |
| | 7 | 2 cm | 1.18 |
| | | 3 cm | 0.69 |
| | | 4 cm | 0.56 |

| **Applied Voltage (V)** | **Receiving Pad Separation** | **Receiving Voltage (dV, V)** |
|---|---|---|
| **3** | **2 cm** | 0.50 |
| | 3 cm | 0.58 |
| | 4 cm | 0.68 |
| 7 | 2 cm | 1.18 |
| | 3 cm | 1.36 |
| | 4 cm | 1.58 |

**SI 5. Multiplexed wireless sensing wearable and implantable.** (A) Wearable "hub" PCB illustrating key components: nRF52832 MCU (BLE communication), ADC converter (sensor data readout), resistive sensing pads, configurable voltage booster for high-voltage pulse actuation, and integrated BLE antenna. (B) PCB fabrication layout highlighting the wearable hub's electronic components and separated ground design. (C) Implantable PCB featuring coin-cell battery, transistor switch, actuator, and receiving pads for triggering via in-body signals. Optional locations for resistors between the transistor base and receiving pad and capacitor between the transistor base and emitter are marked. (D) Android application interface demonstrating real-time monitoring of resistive sensor data and manual pulse stimulation control, providing wireless communication.

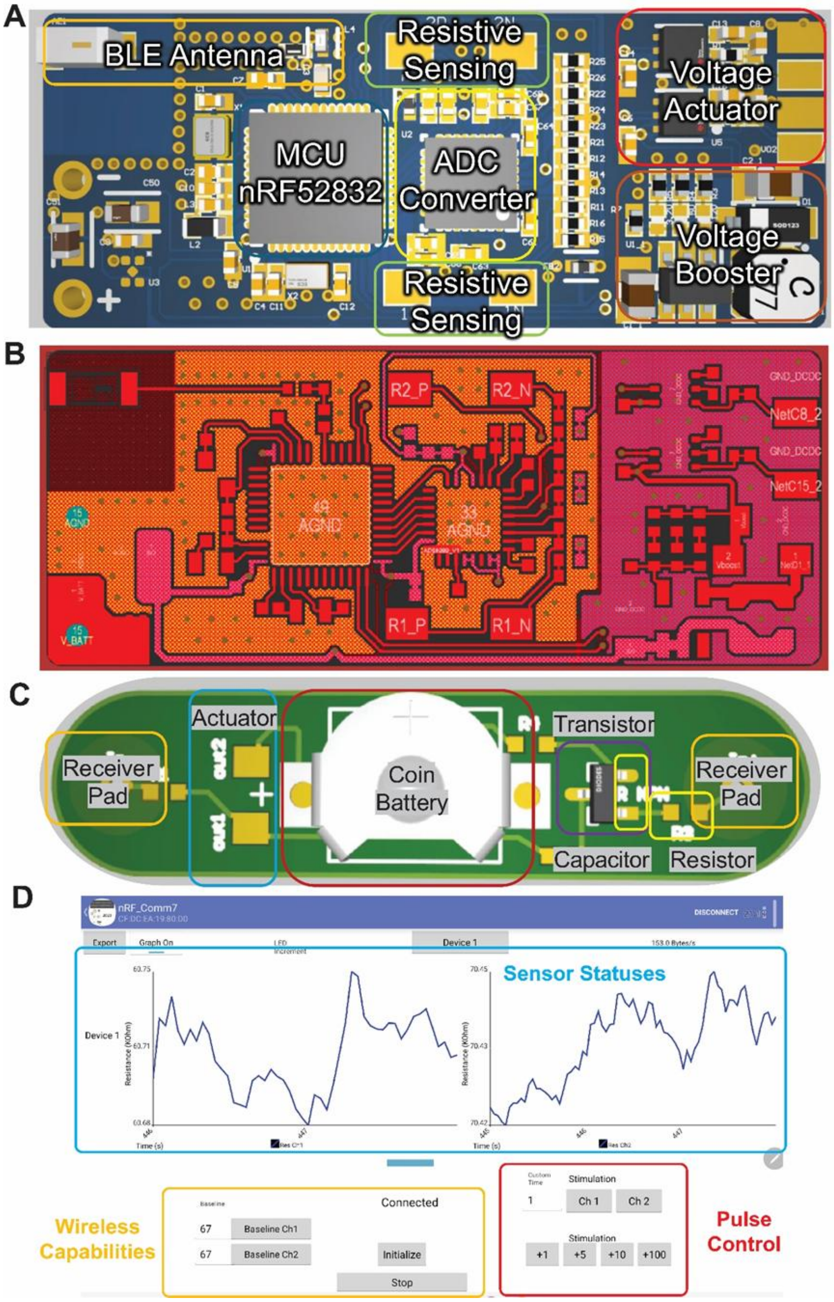

**SI 6. Multiplexed wireless sensing utilizing serpentine traces and laser-induced graphene sensors.** (A) Fabrication of a uniaxial laser-induced graphene (LIG) sensor; Al = aluminum, PDMS = polydimethylsiloxane, Cu = copper, PI = polyimide, SEBS = styrene-ethylene-butylene-styrene block co-polymer. (B) Fabrication of bilayer copper-polyimide serpentine traces. (C) Fabrication of bilayer copper-polyimide interconnect. (D) Strain-resistance calibration curve of 8 LIG sensors (individual = gray lines, average = black line).

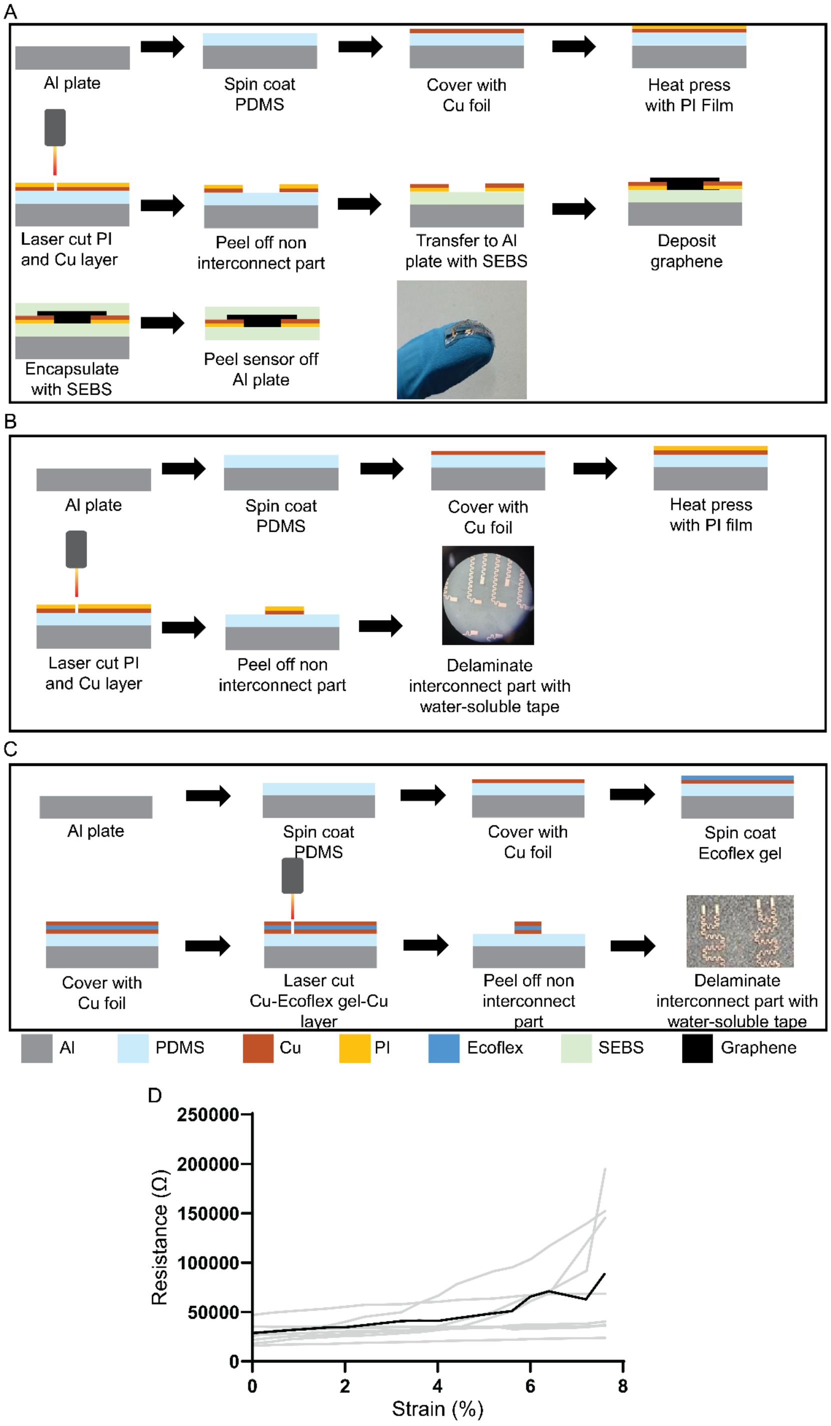

**SI 7. Integrating various actuators for communication via IBC ex vivo.** (A) Ex vivo set up for switching on various electromechanical devices including an LED and a pump. The gate pins of the transistor are connected to tissue-interfacing receiving pads. The electromechanical device is illustrated as a resistive load in the schematic. (B) Utilizing a PNP and an NPN along with a [10 V, 0 V, 1 sec] pulse allows for selective triggering of 2 LEDs. (C) Utilizing a PNP and a MOSFET along with a [6.5 V ,0 V, 3 sec] pulse allows for selective triggering of 2 pumps.

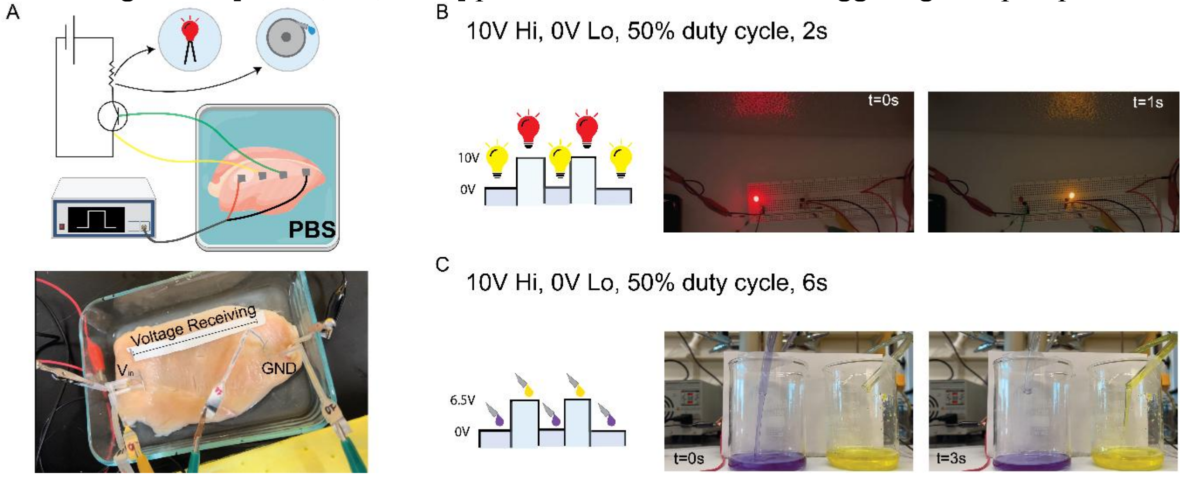
